# The Transcription Factor Bach2 Negatively Regulates Natural Killer Cell Maturation and Function

**DOI:** 10.1101/2022.02.14.480364

**Authors:** Shasha Li, Michael D. Bern, Benpeng Miao, Takeshi Inoue, Sytse J. Piersma, Marco Colonna, Tomohiro Kurosaki, Wayne M. Yokoyama

## Abstract

BTB domain And CNC Homolog 2 (Bach2) is a transcription repressor that actively participates in T and B lymphocyte development, but it is unknown if Bach2 is also involved in the development of innate immune cells, such as natural killer (NK) cells. Here, we followed the expression of Bach2 during NK cell development, finding that it peaked in CD27^+^CD11b^+^ cells and decreased upon further maturation. Bach2 expression positively correlated with that of the transcription factor TCF1 and negatively correlated with genes encoding NK effector molecules as well as genes involved in the cell cycle. Bach2-deficient mice showed increased numbers of terminally differentiated NK cells with increased production of granzymes and cytokines. NK cell-mediated control of tumor metastasis was also augmented in the absence of Bach2. Therefore, Bach2 is a key checkpoint protein regulating NK terminal maturation.

## Introduction

Natural killer (NK) cells are innate lymphoid cells that have spontaneous cytolytic activity against tumor cells and virus-infected cells. The development of NK cells occurs in bone marrow (BM) as well as in secondary lymphoid tissues in both humans and mice. Multipotent hematopoietic stem cells (HSC) give rise to common lymphoid progenitors (CLPs) that can differentiate into all types of lymphocytes. NK cell precursors (NKP) are then derived and later express IL-2R/IL-15R β chain (CD122), defining refined NK precursors (rNKP) (Carotta et al., 2011; Fathman et al., 2011). At this stage in the mouse, commitment to NK cell development occurs, followed by the acquisition of the germline-encoded NK receptors NK1.1, and the cells become immature NK cells. Mature NK cells develop when they gain the expression of DX5 (CD49b), cytotoxic activity, and capacity to produce interferon γ (IFNγ) (Kim et al., 2002). Mature NK cells can be further defined based upon the differential expression of CD27 and CD11b. Starting from double-negative cells being the most immature cells regarding their functionality, the cells upregulate the expression of CD27 then CD11b to become CD27^+^CD11b^-^ (CD27^+^ cells) NK cells then CD27^+^CD11b^+^ (double-positive) NK cells respectively, which undergo homeostatic expansion. Finally, the double-positive cells lose the expression of CD27 and retain expression of CD11b to become terminally differentiated NK cells (CD11b^+^ cells) with increased cytotoxic activity (Chiossone et al., 2009).

The commitment, development, and function of the NK cells are distinctly regulated by multiple transcription factors, which are reflected by the expression of many unique surface markers at different stages of NK cell development. Eomesodermin (Eomes) has been identified as a unique factor required for NK development, distinct from ILC1s which share many similarities in surface makers with NK cells (Gordon et al., 2012; Intlekofer et al., 2005). Kruppel-like factor 2 (KLF2) intrinsically regulates NK cell homeostasis by limiting early-stage NK cell proliferation and guiding them towards trans-presented IL-15 (Rabacal et al., 2016). IRF8 is required for the effector functions of NK cells against viral infection (Adams et al., 2018). T-bet has a broader function in various cells including T cells, ILC1s, and NK cells, and positively regulates the terminal maturation of NK cells (Townsend et al., 2004). Similarly, Zeb2 promotes NK terminal differentiation and may function downstream of T-bet (van Helden et al., 2015). Blimp1 is expressed throughout NK cell maturation and is required for NK homeostasis (Kallies et al., 2011). TCF1 (encoded by *Tcf7* gene) participates in the development of NK cells and its downregulation is required for NK terminal maturation (Jeevan-Raj et al., 2017). A multi-tissue single cell analysis divided NK cells into two major groups based on TCF1 level: high expression of TCF1 correlated with genes expressed in immature NK cells including *Cd27, Xcl1,* and *Kit* while it was inversely correlated with the expression of genes involved in effector function such as *Gzmb, Gzma, Ccl5,* and *Klrg1* (McFarland et al., 2021). Thus, the expression of unique transcription factors at specific developmental stages of the cells appears to generate distinct gene regulatory circuitries. These gene regulatory circuits provide “fingerprints” that may be more reliable to reveal the developmental and functional disparities among phenotypically similar cell populations (Koues et al., 2016). Furthermore, they provide insight into how the development of immune cells, including NK cells, occurs, indicating it is important to clarify the expression pattern and function of unique transcription factors for NK cells during their development.

Identified as a transcriptional repressor, Bach2 forms a heterodimer with Maf family proteins and bind to a DNA motif called T-MARE (TGCTGA G/C TCAGCA), a Maf recognition element to regulate gene expression (Muto et al., 1998). The regulatory function of Bach2 is mediated through its interaction with the super-enhancers (SEs), and its aberrant expression is associated with a variety of autoimmune diseases as well as cancers (Afzali et al., 2017; Marroquí et al., 2014; Roychoudhuri, Eil, et al., 2016). In physiological conditions, Bach2 has been shown to participate in cell development, as previously examined in the adaptive immune system. Bach2 is expressed in CLP and represses genes of myeloid lineage to promote the development of cells in the lymphoid lineage (Itoh-Nakadai et al., 2014). Bach2 was first shown to be a B cell-intrinsic transcription factor that regulates B cell development through inhibiting the expression of Blimp-1 (encoded by *Prdm1* gene) (Muto et al., 1998; Ochiai et al., 2006). The rapid upregulation of Blimp-1 mediated by Bach2-deficiency promotes the terminal differentiation of B cells towards plasma cells even prior to class-switch recombination (CSR) (Muto et al., 2004). Bach2 is also critical in regulating the plasticity of T cells. Under homeostatic conditions, Bach2 maintains T cells in a naïve state, preventing the generation of effector T cells through inhibiting the expression of effector molecules downstream of the TCR signaling (Roychoudhuri, Clever, et al., 2016; Tsukumo et al., 2013). Bach2 expression is reduced during T cell polarization while higher expression of Bach2 in CD4 T cell differentiation promotes the formation of regulatory T cells by repressing genes related to the effector differentiation within helper T cell lineages (Lahmann et al., 2019; Roychoudhuri, Clever, et al., 2016; Roychoudhuri et al., 2013). Regaining the expression of Bach2 after differentiation renders downregulation of pro-inflammatory signals and differentiation into T cell memory cell lineages (Herndler-Brandstetter et al., 2018). Thus, the development and function of the adaptive immune cells are critically controlled by the expression of Bach2 and associated regulatory circuits.

The role of Bach2 in NK cells has not been characterized. Here we found that, at steady state, Bach2 was differentially expressed during NK cell development and terminal maturation. Its deficiency in NK cells resulted in a significantly increased expression of genes involved in NK cytotoxicity. Along with this, NK cells lacking Bach2 expression were more terminally differentiated and demonstrated better control of tumor metastasis. Thus, Bach2 serves as a checkpoint in the terminal maturation of NK cells.

## Results

### Bach2 is expressed at different levels in NK cells at different developmental stages

To characterize the role of Bach2 in NK cell function, we first examined Bach2 expression during different stages of NK development. We used a Bach2^Flag^ reporter mouse in which a 3xFlag tag was fused at the N-terminus of Bach2 protein as described previously (Herndler-Brandstetter et al., 2018). Bach2 expression was detected in common lymphoid progenitors (CLPs). Expression was relatively reduced in pre-NK progenitor cells and refined NK progenitors (rNKp) (Fig. 1A, Supplemental Fig. S1A), reflecting the critical role of Bach2 in the development of lymphoid cells from CLPs (Itoh-Nakadai et al., 2014). During NK specification downstream of CLP, pre-NK cells displayed low levels of Bach2 but regained its expression in refined NK progenitors (Fig. 1A), suggesting Bach2 may play an important role during NK development.

**Fig.1.**
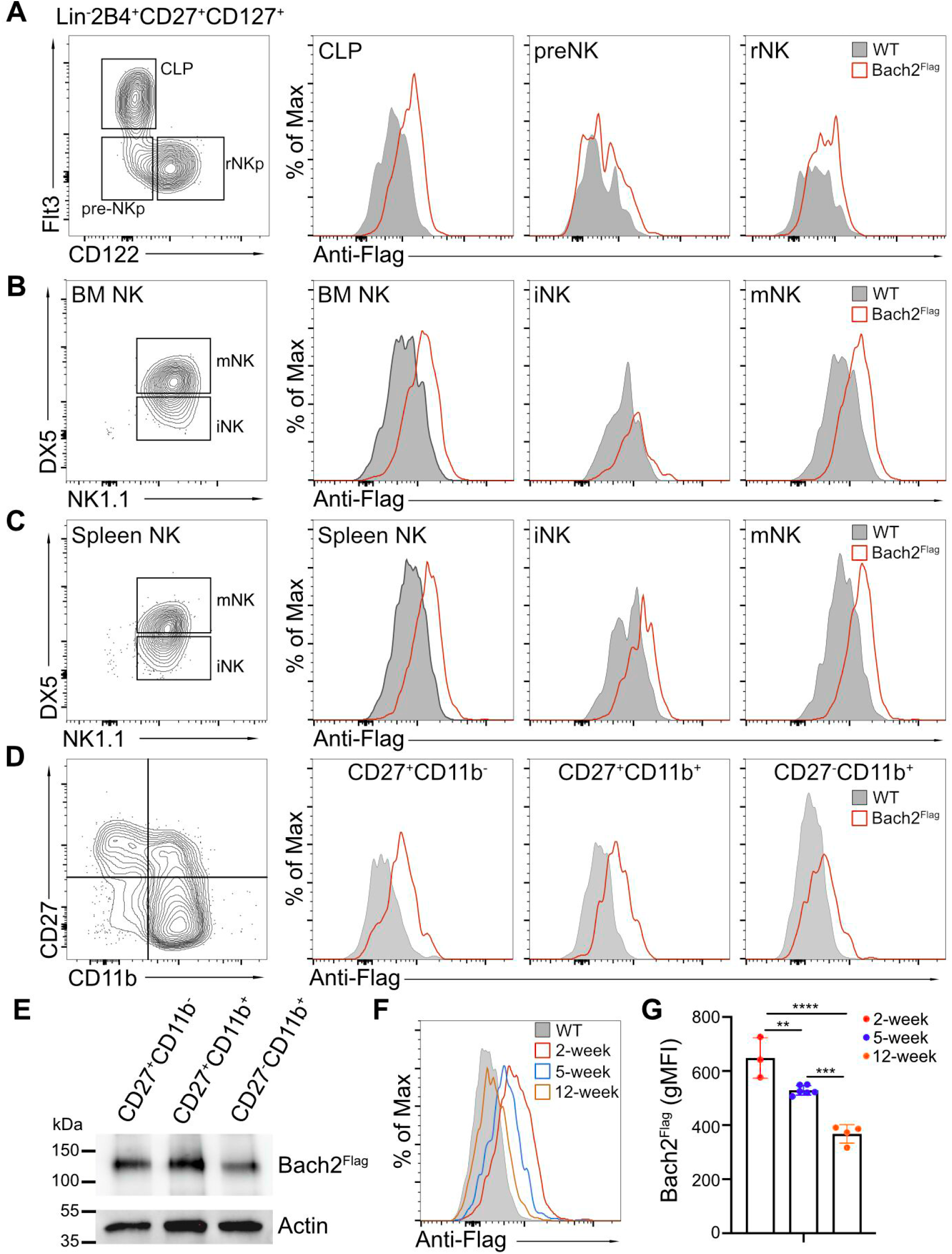
Bach2 expression at different NK cell developmental stages by analysis of Bach2^Flag^ knock-in mouse. (A) BM Lin^-^2B4^+^CD27^+^CD127^+^ cells were subdivided into common lymphoid progenitor (CLP) (Flt3^+^CD122^-^), pre-NK progenitor (pre-NKp) (Flt3^-^CD122^-^), and refined NK progenitor (rNKp) (Flt3^-^CD122^+^). Two individual experiments have been done with a total of four mice per group. (B) CD3^-^CD19^-^CD122^+^ NK cells from BM were subdivided into iNK (DX5^-^NK1.1^+^) and mNK (DX5^+^NK1.1^+^) cells. Bach2 expression was analyzed in the bulk BM NK cells (CD3^-^CD19^-^ NK1.1^+^) and the iNK, mNK subsets. Two individual experiments have been done with a total of four mice per group (C) Same as (B) except cells were harvested from spleen. (D) Splenic NK cells (CD3^-^CD19^-^NK1.1^+^) were subdivided into maturation stages by CD27 and CD11b. Bach2-flag expression was detected in CD27^+^CD11b^-^, CD27^+^CD11b^+^ and CD27^+^CD11b^+^ subsets by flow cytometry. Three individual experiments have been done with a total of four to six mice per group. (E) Splenic NK cells were sorted into CD27^+^CD11b^-^, CD27^+^CD11b^+^ and CD27^-^CD11b^+^ subsets. Bach2-Flag expression in the subsets was detected using Anti-FLAG M2-Peroxidase (HRP) antibody by western blot. Expression of Actin was used as an internal control. Two individual experiments have been done with one mouse each time. (F) Bach2-Flag expression was detected in splenic NK cell (CD3^-^CD19^-^NK1.1^+^) from mice at 2-week (red), 5-week (blue) and 12-week (orange) age. A representative plot was shown for two individual experiments with a total of three to six mice per group. (G) Summary of the geometric MFI (gMFI) of Bach2-Flag on splenic NK cells from mice at indicated age. Data in G are pooled from two independent experiments with a total of three to six mice per group (one-way ANOVA with Tukey’s correction). Histogram overlays show Bach2-Flag expression (open histograms) as compared to cells from wild-type C57BL/6 mice (gray fill). Error bars indicate SD. *\*p* < 0.05; \*\**p* < 0.01; \*\*\**p* < 0.001; \*\*\*\**p* < 0.0001. ns, not significant.

At the CD122^+^CD127^+^ rNKp stage, developing NK cells begin to acquire expression of the NK cell markers, NK1.1 and NKp46 (Fathman et al., 2011). Bach2 was homogenously expressed at higher levels by these NK cells in both the BM and spleen (Fig. 1B and C). At this stage, NK cells further undergo maturation by acquiring the expression of CD49b (DX5). In both the BM and spleen, Bach2 expression can be detected in both immature DX5^−^ and mature DX5^+^ NK cells while mature NK cells have higher Bach2 expression than immature NK cells (Fig. 1B and C, Supplemental Fig. S1B). We further subdivided NK cells by surface expression of CD27 and CD11b. CD27^-^CD11b^-^ and CD27^+^CD11b^-^ subsets (CD27^+^ cells) are regarded as the immature stage. The double-positive CD27^+^CD11b^+^ subset is the intermediate stage, and CD27^-^CD11b^+^ subset (CD11b^+^ cells) represents the most mature stage (Fig. 1D, Supplemental Fig. S1C). The expression of Bach2 was lower in CD27^-^CD11b^+^ cells compared with the other two stages (Fig. 1D). This observation was further confirmed by western blot (Fig. 1E, Supplemental Fig. S1D). Thus, during NK cell terminal differentiation, Bach2 may function mainly in the relatively immature stages.

At the bulk splenic NK cell population level, the expression of Bach2 inversely correlated with the age of the mice: 2-week-old mice had the highest expression, and 15-week-old mice had the lowest expression of Bach2 (Fig. 1F and G). The overall downregulation of Bach2 with age correlated with changes in the maturation profile of NK cells. More than 60 percent of NK cells were immature CD27^+^CD11b^-^ cells in 2-week-old mice while around 40 percent of NK cells in 15-week-old mice were in the most mature stage (CD27^-^CD11b^+^) (Supplemental Fig. S1E). These findings suggest that older mice, having a biased distribution towards mature NK cells, resulted in a reduction of Bach2 levels at the whole population level whereas younger mice displayed the opposite effect.

### Bach2-deficiency drives a transition from immature stem-like phenotype towards mature effector phenotype of NK cells

To further understand the role of Bach2 in NK cell development and function, we generated NK cell-specific Bach2 conditional knockout mice (Bach2^cKO^) by crossing *Bach2*^flox/flox^ mice with *Ncr1*^*i*Cre^ mice in which *Bach2* was specifically deleted in NKp46-expressing cells which mainly include NK cells among splenocytes. As a control, *Bach2*^flox/flox^ mice (referred to as control mice) were used. NK cell number in Bach2 knockout mice is unchanged compared to control mice (Supplemental Fig. S2A). We assessed the expression of the Ly49 and NKG2A/CD94 receptors, important for missing-self mediated cytotoxicity, in splenic NK cells from Bach2^cKO^ and control mice. We detected only slight changes in the expression profile of the repertoire of these receptors with Bach2-deficiency while CD94 is significantly downregulated (Supplemental Fig. S2B).

We performed RNA-seq analysis compared between Bach2-deficient NK cells and Bach2-sufficient NK cells. We sorted the splenic NK cells from Bach2^cKO^ and control mice to perform the RNA-seq analysis (Supplemental Fig. S2C). Principal component analysis (PCA) revealed significant changes transcriptionally in NK cells at the quiescent condition with Bach2-deficiency (Fig. S2D). We found 133 genes downregulated and 210 genes upregulated in Bach2^cKO^ NK cells as compared to control NK cells (Bach2^cKO^ versus control) (Fig. 2A and Supplemental Table S1). The transcripts with decreased expression corresponded to genes involved in T cell differentiation, cell development, and cell homeostasis pathways (Fig. 2A). These gene signatures suggested Bach2 controlled NK cell differentiation. Specifically, the top 2 downregulated genes included *Kit* and *Tcf7,* which were previously shown to be responsible to maintain the stemness of the T cells (Siddiqui et al., 2019). Regarding to NK cells, the loss of *Tcf7* expression led to enhanced NK cell terminal maturation (Jeevan-Raj et al., 2017). We also detected the downregulation of *Cd27, Ccr7,* and *Cd69.* (Fig. 2B). The downregulation of *Ccr7* was also previously shown to be correlated with human NK cell differentiation towards effector phenotype (Hong et al., 2012). On the other hand, the transcripts with an elevated expression included the genes involved in the cell cycle, cell proliferation, and inflammatory response pathways (Fig. 2A), suggesting a skewing towards an effector phenotype as a result of Bach2-deficiency. Indeed, among the upregulated genes were those involved in NK cell effector functions such as *Klrg1, Gzmb, Gzmk,* and *Ccl5* (Fig. 2B) (Bezman et al., 2012), indicating that lack of Bach2 expression facilitated the differentiation of NK cells towards terminal maturation.

**Fig. 2.**
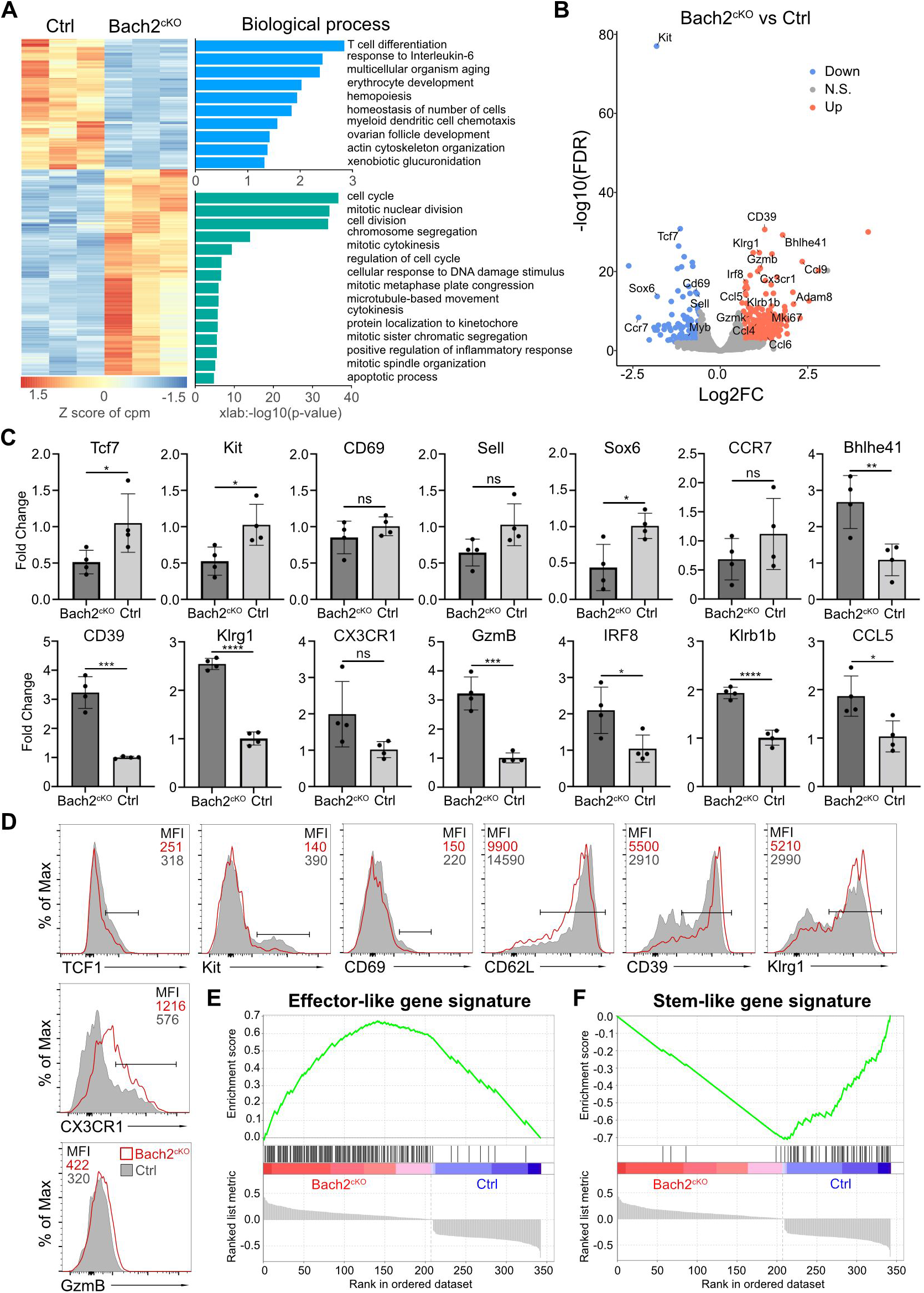
RNA-seq analysis reveals Bach2-deficiency in NK cells promotes the terminal maturation of NK cells with elevated effector function. (A) Heatmap of differentially expressed genes in NK cells compared between control and Bach2^cKO^ mice in RNA-seq analysis. Each column represents total splenic CD3^-^CD19^-^ NK1.1^+^NKp46^+^ cells from an individual mouse. Cells were sorted from two individual experiments with four mice per group. The data were analyzed with the Database for Annotation, Visualization and Integrated Discovery (DAVID) Gene Ontology (GO) analysis for the biological process using the genes differentially expressed from NK cells between control and Bach2^cKO^ mice. (B) Volcano plot shows the differential gene expression between control and Bach2^cKO^ splenic NK cells. Highlighted are genes discussed in the text. (C) Quantitative real-time PCR (qPCR) validation of selected genes. Data are shown with four mice per group from two individual experiments (student’s *t* test). (D) Flow cytometry plots of indicated protein expression in Bach2^cKO^ mice (open histograms) or control mice (gray fill). Two experiments have been done with a total of two mice per group. (E and F) GSEA illustrating the enrichment of effector-like (E) and stem-like (F) gene signatures in Bach2^cKO^ and control splenic NK cells. Error bars indicate SD. *\*p* < 0.05; \*\**p* < 0.01. ns, not significant.

We confirmed the transcriptomics data by quantitative PCR (qPCR) analysis performed on sorted splenic NK cells from the Bach2^cKO^ and control mice respectively (Fig. 2C). We detected a significantly lower expression of genes for example *Tcf7, Kit,* and *Sox6* in Bach2 deficient NK cells. For other genes such as *Cd69, Cd62l (Sell),* and *Ccr7,* we did not observe significant differences but they all had a trend of downregulation in NK cells lacking Bach2 expression. The genes we picked with upregulation in RNA-seq data were confirmed to be increased by qPCR analysis. These genes included *Gzmb, Klrg1, Ccl5, Klrb1b, Cd39,* etc. except for *Cx3cr1* which did not reach significance even though displaying a trend of upregulation. Consistent with their transcription level, the protein encoded by the genes was also revealed to be changed caused by Bach2-deficiency (Fig. 2D). TCF1, Kit, CD69, and CD62L were all shown to be decreased at the protein level. Although *Tcf7* had a dramatic decrease transcriptionally, its protein was only slightly downregulated. On the other hand, CD39, KLRG1, CX3CR1, and Granzyme B were elevated followed by their changes at the RNA level. Thus, we confirmed that our RNA-seq data were reliable to reflect the impact of Bach2 in regulating the expression of various genes in NK cells.

Next, we asked how Bach2 participated in NK cell biology. It was reported that the enforced Bach2 expression in exhausted CD8^+^ T cells resulted in the cells becoming exclusively stem-like precursor exhausted CD8^+^ T cells, preventing their further differentiation into terminal exhausted CD8^+^ T cells (Yao et al., 2021). We used gene-set enrichment analysis (GSEA) to determine the effect of Bach2-deficiency by comparing against the gene signatures of stem-like CD8^+^ T cells and terminally differentiated effector-like CD8^+^ T cells. We found genes upregulated in NK cells induced by Bach2-deficiency positively correlated with terminal differentiated effector-like gene signatures (Fig. 2E) whereas the genes downregulated showed a stem-like signature (Fig. 2F). Another comparison was performed using GSEA analysis and we found Bach2-sufficient NK cells displayed a naïve CD8^+^ T cell signature (Fig. S2E) while Bach2-deficient NK cells resembled activated effector CD8^+^ T cells (Fig. S2F). These data suggested that Bach2 expression suppressed terminal differentiation of NK cells by repressing many effector genes.

### Bach2 restrained terminal maturation of NK cells

Based on the expression pattern of Bach2 in NK cells as well as RNA-seq and confirmatory qPCR data in Bach2-deficient cells, we tested whether Bach2 indeed restricted the terminal maturation of the immature NK cells. To address this, we analyzed the maturation profile of the NK cells compared between Bach2^cKO^ and control mice. In the bone marrow, we did not observe a significant difference between control and Bach2^cKO^ mice regarding the frequency of the CD27^+^ cells or CD11b^+^ cells (Fig. 3A). However, an altered maturation profile of NK cells was detected in the spleens of Bach2^cKO^ mice, i.e., there were more cells with mature NK cell phenotype (CD11b^+^) and fewer NK cells at the immature DP stage as compared to control mice (Fig. 3B). Consistent with this pattern, expression of KLRG1, a marker of terminally mature splenic NK cells was also increased at the population level, and we detected the percentage of KLRG1^+^ NK cells dramatically increased in Bach2^cKO^ mice (Fig. 3C). To confirm the lack of influence of T cells and B cells on NK cell development and maturation, we evaluated NK cells in germline Bach2-deficient mice on the *Rag1^-/-^* background. The maturation of NK cells was analyzed in both *Bach2^-/-^ Rag1^-/-^* mice and *Rag1^-/-^* mice. Similar to the results presented in Bach2^cKO^ mice, we found the frequency of CD27^+^ NK cells, specifically the DP (CD27^+^CD11b^+^) NK cells, was significantly reduced while the frequency of CD11b^+^ NK cells was markedly increased in both bone marrow (Fig. 3D) and spleen (Fig. 3E) in *Bach2^-/-^* mice. In agreement, the expression of KLRG1 was upregulated and we also found more cells (~80%) expressing KLRG1 in Bach2-deficient mice compared to Bach2-sufficient mice (~50%) (Fig. 3F). Taken together, NK cells were skewed towards the most mature NK cells in Bach2-deficient mice as compared to Bach2-sufficient mice, with a concomitant decrease in the immature NK cells.

**Fig. 3.**
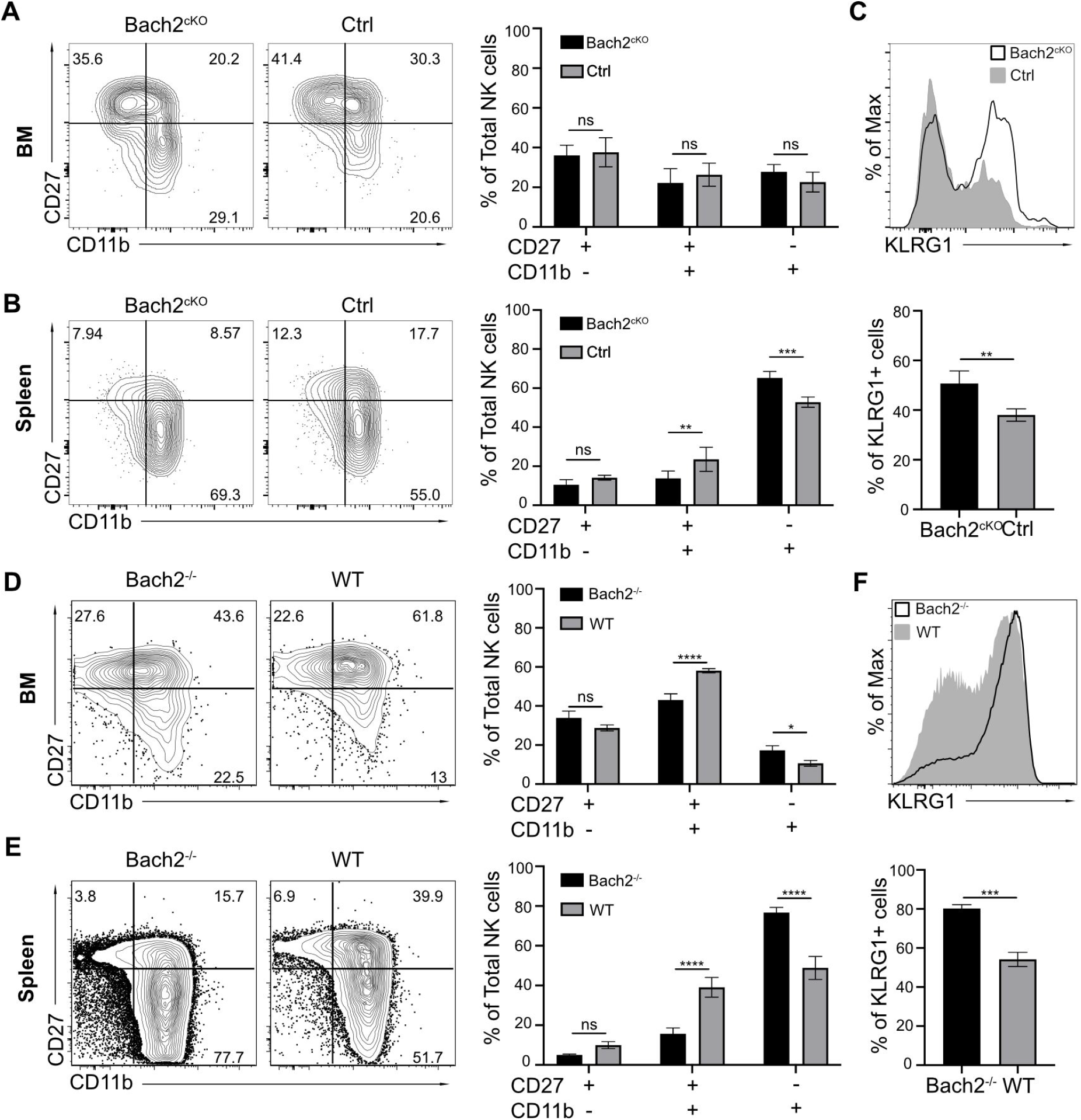
Bach2-deficiency increases NK cells with terminally differentiated phenotype. (A and B) Representative flow cytometry plots of NK cells (CD3^-^CD19^-^NK1.1^+^NKp46^+^) separated into maturation stages by CD27 and CD11b expression from BM (A) or spleen (B) in control mice and Bach2^cKO^ mice. Percentage of different subsets were plotted. Data were pooled from three independent experiments with a total of four to six mice per group (two-way ANOVA with Bonferroni correction). (C) Representative histogram of KLRG1 expression on splenic NK cells (CD3^-^CD19^-^NK1.1^+^NKp46^+^) from control mice and Bach2^cKO^ mice. Percentage of NK cells that express KLRG1 from control mice and Bach2^cKO^ mice were pooled from three independent experiments with a total of four to six mice per group (student’s *t* test). (D and E) Representative flow cytometry plots of total NK cells (CD3^-^CD19^-^NK1.1^+^NKp46^+^) maturation stages separated by CD27 and CD11b expression from BM (D) or spleen (E) in Rag1^-/-^Bach2^-/-^ (Bach2^-/-^) and Rag1^-/-^ (WT) mice. Percentage of different subsets were plotted. Data were shown for three mice per group from one experiment (two-way ANOVA with Bonferroni correction). (F) Representative histogram of KLRG1 expression on splenic NK cells (CD3^-^CD19^-^ NK1.1^+^NKp46^+^) from Rag1^-/-^Bach2^-/-^ (Bach2^-/-^) and Rag1^-/-^ (WT) mice. The percentage of NK cells that express KLRG1 was shown for three mice per group from one experiment (student’s *t* test). Error bars indicate SD. *\*p* < 0.05; \*\**p* < 0.01; \*\*\**p* < 0.001; \*\*\*\**p* < 0.0001. ns, not significant.

### B16 tumor growth and metastasis is controlled by NK cells with Bach2-deficiency

Since differential gene analysis showed many effector molecules, especially cytotoxic genes were increased with Bach2-deficiency, we evaluated whether Bach2-deficiency resulted in changes in the rejection of target cells. We asked whether the increased representation of more mature NK cells due to Bach2-deficiency would result in better control of tumor metastases *in vivo.* We assessed the role of Bach2 in B16F10 metastasis. In tumor metastasis studies, 2.5×10^5^ B16F10 cells were intravenously injected into Bach2^cKO^ or control mice and tumor metastases were evaluated two weeks later by counting the black colonies formed in lungs (Fig. 4A). The lung metastases of B16F10 tumors in Bach2^cKO^ mice were dramatically reduced to 25% of metastatic colonies found in control mice (Fig. 4B). This control of tumor metastases was NK cell-dependent because both mice displayed a higher and similar number of metastatic tumor colonies when NK cells were depleted with anti-NK1.1 (PK136) one day prior to injection (Fig. 4B). In summary, NK cells in Bach2-deficient mice are more efficient in controlling tumor progression and metastasis.

**Fig. 4.**
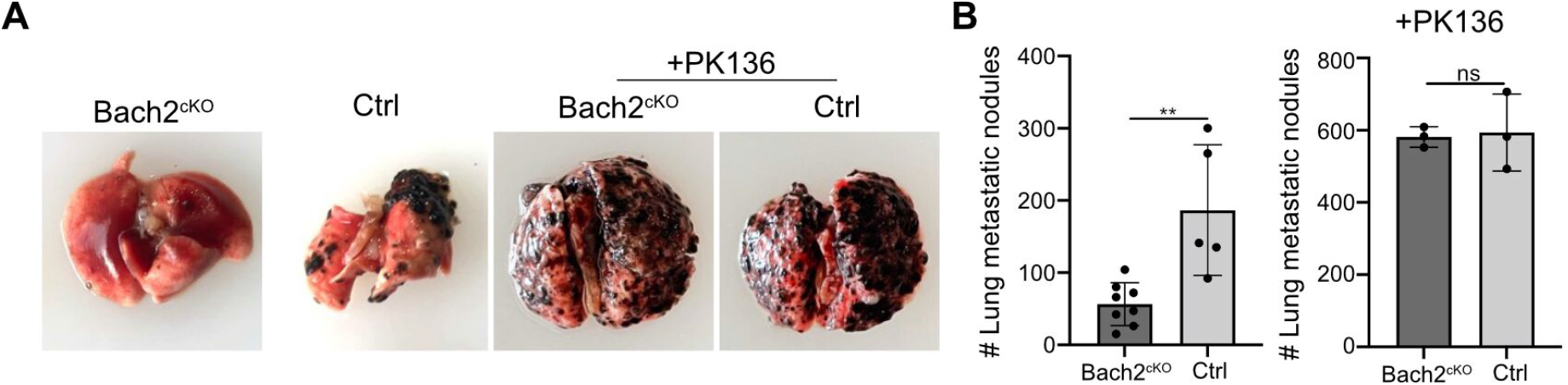
Lack of Bach2 expression in NK cells suppresses B16F10 tumor metastasis and growth. (A) Representative picture of lung metastatic nodules in Bach2^cKO^ mice and control mice under steady state or anti-NK1.1 (PK136) depletion. (B) The number of B16F10 metastatic nodules in lung from Bach2^cKO^ and control mice with or without anti-NK1.1 (PK136) depletion. Data were pooled from three independent experiments with a total of three to eight mice per group (student’s *t* test). Error bars indicate SD. *\*\*p* < 0.01.

## Discussion

Here we found that Bach2 is highly expressed in functional immature NK cells (CD27^+^ NK cells) and gradually downregulates its expression at the terminal stage of NK maturation (CD11b^+^ NK cells). In line with this, we demonstrated that Bach2-deficiency caused a biased NK cell development towards terminal differentiation in an NK cell intrinsic manner. Bach2 deficient NK cells displayed increased cytotoxic gene expression and were more potent in controlling tumor metastases.

Our prior knowledge of Bach2 was largely from other cells in the lymphoid lineage such as B cells and T cells. Bach2 played a critical role in the development of the lymphoid cells: its presence in common lymphoid progenitor (CLP) repressed the expression of genes important for myeloid cells, promoting the development of T, B cells (Itoh-Nakadai et al., 2014). However, in bone marrow, the expression of Bach2 increased after commitment of stem cells to the B cell lineage with its expression high in pre-pro B cells, pro-B cells, pre-B cells, and immature B cells. In B cells (Muto et al., 1998), Bach2 served as a checkpoint protein that inhibited the expression of the immunoglobulin heavy chain of activated p53 by competing with BCL6 for functional VDJ rearrangements (Muto et al., 1998; Swaminathan et al., 2013). Bach2 also suppressed the differentiation of activated B cells to plasma cells by inhibiting the expression of the Blimp-1 (encoded by *Prdm1* gene), which allowed the class switch recombination (CSR) and somatic hypermutation (SHM) to take place before becoming plasma cells abruptly (Kometani et al., 2013; Muto et al., 2004; Ochiai et al., 2006). Here, we also detected a relatively high expression of Bach2 in CLP but there was a reduction in pre-NK progenitors and refined NK progenitors. We showed Bach2 started to gain its expression in NK cells after the acquisition of germline-encoded NK receptors such as NK1.1 and NKp46, a stage when NK cells displayed an effector program. The differential expression pattern of Bach2 in the early stage between NK cell development and B cell development indicated a divergent trajectory of CLP for the commitment of B cells or NK cells.

Bach2 was specifically highly expressed in CD27^+^ cells but not in CD11b^+^ terminal differentiated cells. Interestingly, maturing NK cells upregulate CD27 transiently followed by the upregulation of CD11b and KLRG1 (Chiossone et al., 2009; Huntington et al., 2007). During this transition, NK cells lose their homeostatic expansion capacity but acquire cytotoxic activity (Chiossone et al., 2009; Huntington et al., 2007), suggesting the main role of Bach2 is to maintain the homeostasis of the most mature NK cells. Indeed, when we examined the differential gene expression in the context of Bach2-deficiency specifically in mature NK cells, we detected an upregulation of a series of genes related to cell proliferation, immune effector molecules, and cell apoptosis. In contrast, genes associated with cell development and homeostasis were downregulated with Bach2-deficiency, suggesting that Bach2 might be required for NK cell self-renewal at steady state.

The genes we detected with differential expression patterns correlated very well with the gene signatures observed in CD8 T cells, their cytotoxic counterparts in the adaptive immune system. One of the genes, *Tcf7,* was particularly interesting and may be important for the mechanism of the regulation of NK development by Bach2. *Tcf7* (encoding TCF1) is highly expressed in naïve T cells, decreased in effector T cells, and regained its expression in memory cells, showing its role in the maintaining pluripotency of the T cells (Willinger et al., 2006; Zhao et al., 2010).

Similarly, Bach2 also maintained T cells in a naïve status under homeostatic conditions, preventing the generation of effector T cells through inhibiting the expression of effector molecules downstream of TCR signaling (Roychoudhuri, Clever, et al., 2016; Tsukumo et al., 2013). On the other hand, TCF1 was recently shown to be a hallmark of stem-like precursor exhausted CD8 T cells with self-renewal capability and can differentiate into terminal effector-like exhausted CD8 T cells which lacked TCF1 expression (Utzschneider et al., 2020). It was shown Bach2 also played a positive role in maintaining the pool of these stem-like precursors exhausted CD8 T cells as enforced overexpression of Bach2 resulted in the cells retaining this stem-like condition while knockout of Bach2 led to terminal differentiation of the cells (Yao et al., 2021). Our data also showed the link between Bach2 and TCF1: Bach2-deficiency caused downregulation of TCF1 transcription. More importantly, TCF1 has been previously shown to participate in NK development and its downregulation was required for NK cell terminal maturation (Jeevan-Raj et al., 2017). It would be important to understand whether Bach2 and TCF1 will have some interaction in regulating NK cell development. Given that Bach2 is a transcriptional repressor through its interaction with the super-enhancers (SEs), it is possible that Bach2 may directly regulate TCF1 expression to impact NK maturation. However, since we only detected a minor decrease of TCF1 at the protein level in Bach2-deficient NK cells, to what extent this regulation would impact NK development via TCF1 requires further investigation.

Human NK cells encompass two major subsets, known as CD56^dim^ and CD56^bright^ NK cells. CD62L (encoded by *Sell)* and CCR7 were shown to be highly expressed by CD56^bright^ NK cells and drove their migration to secondary lymphoid tissues (Campbell et al., 2001; Frey et al., 1998). CD56^bright^ NK cells also expressed high levels of c-Kit for their homeostatic proliferation (Matos et al., 1993). In agreement with this, *Sell*, *Ccr7,* and *Kit* genes were all downregulated in Bach2-deficient NK cells in our data in mice. In contrast, CD56^dim^ NK cells displayed a high density of CX3CR1 for the migration to tissues and higher cytotoxic activity by increased expression of perforin and various granzymes (Campbell et al., 2001), which resembled our Bach2-deficient NK phenotypes in mice. More importantly, Bach2 has been demonstrated to be highly expressed by CD56^bright^ NK cells and with low expression in CD56^dim^ NK cells (Holmes et al., 2021). Another study of regulome analysis in human NK proposed that Bach2-mediated gene suppression relied upon inhibiting BLIMP1 (encoded by *PRDM1* gene) expression, and BLIMP1 repressed the TCF1-LEF1-MYC-induced homeostatic expansion of NK cells (Koues et al., 2016). However, we did not detect a significant upregulation of the *Prdm1* gene in Bach2 knockout NK cells in our data. Therefore, the function of Bach2 may be conserved between human and mouse for its regulatory circuitries but still needs further exploration.

NK cells are currently being studied in clinical trials as potential targets for cancer immunotherapy. Our study shows that, in mice, Bach2 functions as a checkpoint to restrain NK cell cytotoxicity and Bach2-deficiency leads to enhanced NK cell-mediated control of B16 melanoma metastases. Studies in humans have also suggested that Bach2 may play a similar role in human NK cells (Koues et al., 2016). As a result, our study suggests that Bach2 may be a novel target for checkpoint inhibition of NK cells for cancer immunotherapy.

## Materials and Methods

### Mice

Wild-type C57BL/6 (B6) mice and RAG1^-/-^ mice were purchased from The Jackson Laboratories. Bach2^Flag^ knock-in mice have been described before (Herndler-Brandstetter et al., 2018) as have Bach2^flox/flox^ mice (Kometani et al., 2013). NK cell Bach2 conditional knockout mice were generated by crossing Bach2^flox/flox^ mice with Ncr1^iCre^ mice (Narni-Mancinelli et al., 2011) from Eric Vivier (CNRS-INSERM-Universite de la Mediterranee, Marseille, France). ES cells for Bach2^-/-^ (Bach2^tm1e^) mice were purchased from the EuComm program. The mice were derived from ES clone EPD0689_1_H03, ES line JM8A3.N1. Animal experiments were performed with 6- to 12-week male or female mice, except for those specifically indicated. Mouse studies were conducted in accordance with the institutional ethical guidelines through institutional animal care and use committee (IACUC) protocol that was approved by the Animal Studies Committee of Washington University (#20180293).

### Antibodies and Flow Cytometry

The following antibodies and reagents were purchased from eBioscience: anti-CD127 (A7R34), anti-CD3e (145-2C11), anti-CD19 (eBio1D3), anti-CD49b (DX5), anti-NK1.1 (clone PK136), anti-CD27 (LG.TF9), anti-CD11b (M1/70), anti-NKp46 (29A1.4), anti-CD39 (24DMS1), anti-Granzyme B (NGZB), anti-Ly49A (eBio12A8), anti-Ly49D (eBio4E5), anti-Ly49EF (CM4), anti-Ly49I (YLI-90), anti-CD94 (18d3), anti-NKG2A (16a11),anti-TER-119 (TER-119), Fixable Viability Dye eFluor 506. The following antibodies were purchased from BD Biosciences: anti-CD244.2 (2B4), anti-TCF-7/TCF-1 (S33-966), anti-Ly49F (HBF-719), anti-Ly6-G and Ly-6C (RB6-8C5), anti-CD122 (TM-β1), anti-CD69 (H1.2F3), anti-CD62L (MEL-14), anti-Ly-49G2 (4D11), anti-CD135 (A2F10.1). The following antibodies were purchased from Biolegend: anti-DYKDDDDK tag (L5), anti-KLRG1 (MAFA) (2F1/KLRG1), anti-CD117 (2B8), anti-CX3CR1 (SA011F11). Anti-Ly49H (3D10) and anti-Ly49C (4LO33) were produced in-house. BM or spleen cells were treated with RBC lysis buffer to remove erythrocytes. Then cells were treated with 2.4G2 (anti-Fc RII/III) hybridoma supernatants to block Fc receptors. Surface staining was performed on ice in FACS staining buffer (1% BSA, 0.01% NaN3 in PBS). The labeling of BM progenitor populations has been described before (Jeevan-Raj et al., 2017). Lineage positive cells were labeled by a cocktail of biotin-conjugated anti-CD3e, CD19, NK1.1, CD11b, Gr-1, Ter-119 antibodies. The resulting lineage-negative cells (Lin^-^) were further stained to identify CLP, pre-NK progenitor, and rNK progenitor as indicated. For intracellular staining of Bach2^Flag^, TCF1, and GzmB, the Foxp3 transcription factor staining buffer set (eBioscience) was used according to the manufacturer’s protocols. Samples were collected by FACS Canto (BD Bioscience), and data were analyzed by FlowJo.

### Western Blot

NK cells from spleen were enriched by the EasySep Mouse NK cell isolation kit (STEMCELL Technologies) according to the manufacturer’s instructions. Enriched NK cells were then labeled by indicated surfaced markers and sorted into different subsets. Sorted cells were lysed in RIPA buffer in the presence of Halt Protease Inhibitor Cocktail (Thermo Scientific, 78429) on ice. Lysates were denatured in 2x Laemmli sample buffer (Bio-Rad) and resolved by SDS-PAGE. Proteins were transferred to NC membrane and probed with indicated antibodies. Anti-FLAG M2-Peroxidase (HRP) (A8592) was purchased from Sigma. Beta-Actin Rabbit antibody (4967S) and anti-rabbit IgG HRP-linked antibody (7074S) were purchased from Cell signaling.

### RNA-sequencing and quantitative PCR

NK cells from the spleen were enriched by the EasySep Mouse NK cell isolation kit (STEMCELL Technologies) according to the manufacturer’s instructions. Enriched cells were then sorted into CD3^-^CD19^-^NK1.1^+^NKp46^+^ cells. RNA was purified by PureLink RNA Mini Kit (Ambion, 12183018A). RNA-sequencing was performed by the Genome Technology Access Center at Washington University School of Medicine. NovaSeq 6000 was used for sequencing. RNA-seq reads were then aligned to the Ensembl release 76 primary assembly with STAR version 2.5.1a (Dobin et al., 2013). Gene counts were derived from the number of uniquely aligned unambiguous reads by Subread/featureCount version 1.4.6-p5 (Liao et al., 2014). Low expressing genes were filtered with the criteria of cmp>1 in at least three samples. Two outliers were removed. RUVr method (k=1) in RUVseq R package was used to remove batch effect. Differential gene expression was determined using the EdgeR R package with FDR<0.01 and log2 fold change > log2(1.5) as the thresholds. Heatmaps were generated with pheatmap R package. Principal components analysis (PCA) was performed by prcomp function of R. Gene set enrichment pathways analysis was done using the Broad Institute’s GSEA software by comparing signature databases from GSE83978 and GSE77857. The data in this paper have been uploaded to Gene Expression Omnibus under accession number GSE196530 that also includes Supplemental Table S1.

For qPCR, cDNA was synthesized by ProtoScript II Reverse Transcriptase (NEB, M0368S). Pre-designed primers for indicated genes were obtained from IDT. Quantitative real-time PCR was performed by PowerUp SYBR Green Master Mix Kit (Fisher Scientific) on a StepOnePlus real-time PCR system (Thermo Fisher Scientific). Relative gene expression was normalized to betaactin and calculated by the ΔΔCt method.

### B16F10 Metastasis assay

B16F10 cells were maintained in R10 (RPMI 1640 medium (Gibco) containing 10% FBS, 1% Penicillin/Streptomycin, 1% L-glutamine, 55μM 2-mercaptoethanol). Before injection, 2.5×10^5^ B16F10 cells were resuspended in 300ul PBS, and intravenously injected into mice. After 14 days, tumor metastasis was evaluated by counting the black colonies formed in the lung under a dissecting microscope. The blind analysis is performed for counting.

### Statistical analysis

Statistical analyses were performed using GraphPad Prism 9 software. The statistical test used is stated in the figure legend. Data are presented as mean ± SD as stated in the figure legend. Statistical significance was determined as indicated. p < 0.05 was considered statistically significant.

## Acknowledgments

We thank Susan Gilfillan for helping with the establishment of the Bach2^-/-^ (Bach2^tm1e^) cell. We thank Beatrice Plougastel-Douglas and Anna Sliz for helpful discussions.

## Supplementary Figure Legends

**Fig. S1.**
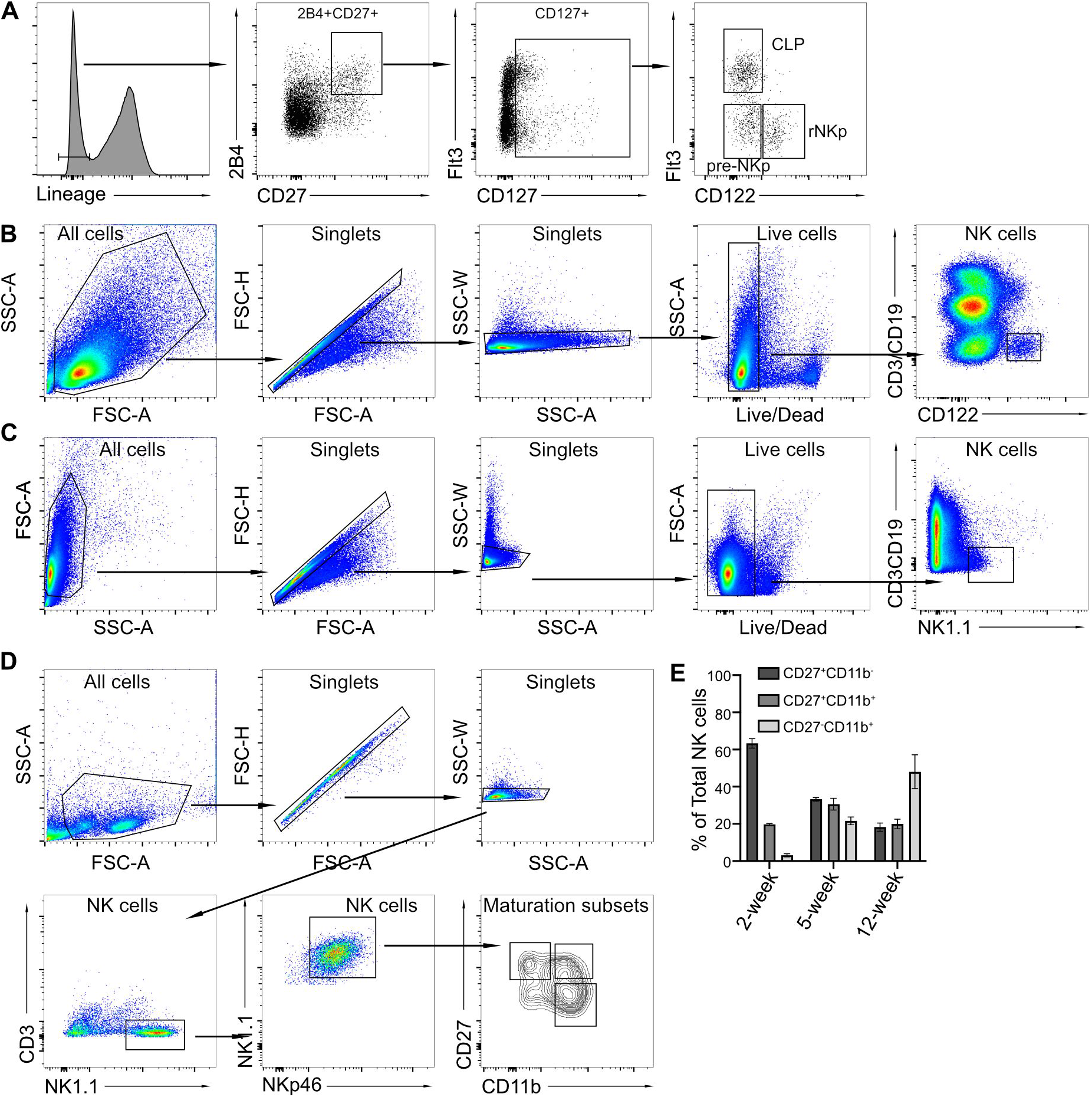
Gating strategy for flow cytometry analysis. (A) Representative flow cytometry gating strategy showing the common lymphoid progenitor (CLP) cells, pre-NK progenitors, and refined NK progenitor cells in BM. (B) Representative flow cytometry gating strategy for immature and mature NK cells from bone marrow and spleen, corresponding to Figs. 1B and C. (C) Representative flow cytometry gating for NK cells, corresponding to Fig. 1D. (D) Representative flow cytometry for the sorting of different subsets of NK cells in the spleen, corresponding to Fig. 1E. (E) Maturation stages of splenic NK cells separated by CD27 and CD11b expression from mice at the indicated age. Data are shown for three mice per group from one experiment.

**Fig. S2.**
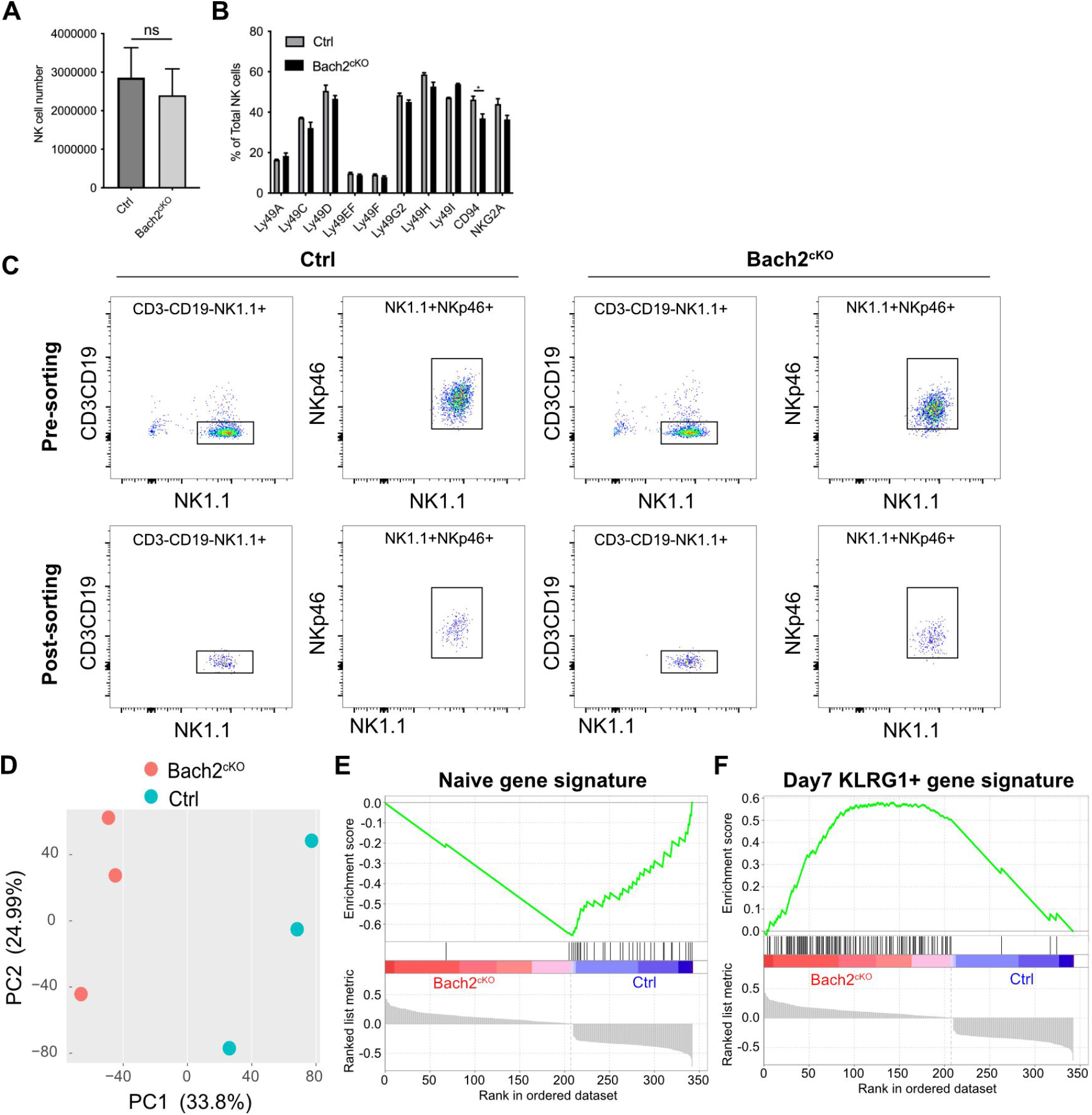
Bach2-deficiency in NK cells resembles activated effector CD8^+^ T cells. (A) Total number of NK cells from the spleen of Bach2^cKO^ or control mice. Data are shown for three mice per group from one experiment (student’s *t* test). ns, not significant. (B) The expression profile of various NK receptors (% of total NK cells) in control and Bach2^cKO^ mice. Data were pooled from two independent experiments with a total of four to six mice per group (two-way ANOVA with Bonferroni correction). \**p* < 0.05. (C) Flow cytometry sorting of splenic NK cells (CD3^-^CD19^-^NK1.1^+^NKp46^+^) from Bach2^cKO^ or control mice for RNA-seq analysis (Fig. 2A). Post-sorting flow cytometry shows the purity of the NK cells. (D) Principal component analysis (PCA) of differentially expressed genes (log2FC>log2(1.5) and FDR<0.01 or log2FC<-log2(1.5) and FDR<0.01) for splenic NK cells isolated from control and Bach2^cKO^ mice. (E and F) GSEA illustrating the enrichment of naïve (E) and day7 Klrg1-positive (F) gene signatures in Bach2^cKO^ and control splenic NK cells.

**Supplemental Table S1** Differential gene expression in NK cells between Bach2^cKO^ and control mice.

**Figure 1-source data.** Bach2 expression in different subsets by western blot. Splenic NK cells were enriched from splenocytes of Bach2^Flag^ mice. Enriched NK cells (CD3^-^ NK1.1^+^) from Bach2^Flag^ mice were further sorted into CD27^+^CD11b^-^, CD27^+^CD11b^+^ and CD27^-^ CD11b^+^ subsets. (A) Bach2 expression in the subsets was detected using Anti-FLAG M2-Peroxidase (HRP) antibody by western blot. (B) Expression of Actin was used as an internal control. Splenocytes from WT mice were used as negative control. Splenocytes from Bach2^Flag^ mice were used as positive control. Two individual experiments have been done with one mouse each time. (C) Composite figure using source data from A and B.

